# Response-locked theta dissociations reveal potential feedback signal following successful retrieval

**DOI:** 10.1101/2024.01.11.575166

**Authors:** Devyn E. Smith, Justin R. Wheelock, Nicole M. Long

**Author notes:** Corresponding Author: Nicole Long; twitter: @dorsolateralpfc).

## Abstract

Successful memory retrieval relies on memory processes to access an internal representation and decision processes to evaluate and respond to the accessed representation, both of which are supported by fluctuations in theta (4-8Hz) activity. However, the extent to which decision making processes are engaged following a memory response is unclear. Here, we recorded scalp electroencephalography (EEG) while human participants performed a recognition memory task. We focused on response-locked data, allowing us to investigate the processes that occur prior to and following a memory response. We replicate previous work and find that prior to a memory response theta power is greater for identification of previously studied items (hits) relative to rejection of novel lures (correct rejections; CRs). Following the memory response, the theta power dissociation ‘flips’ whereby theta power is greater for CRs relative to hits. We find that the post-response ‘flip’ is more robust for hits that are committed quickly, potentially reflecting a positive feedback signal for strongly remembered experiences. Our findings suggest that there are potentially distinct processes occurring before and after a memory response that are modulated by successful memory retrieval.

## 1 Introduction

Successful remembering is dependent on both memory processes to access stored representations and decision processes to evaluate and respond to the accessed representation. How these processes unfold over time, and the underlying neural mechanisms, are critically important to memory success, yet our understanding of these processes remains limited. In particular, convergent evidence from scalp electroencephalography (EEG) studies in both the memory literature (Klimesch, Doppelmayr, Schimke, & Ripper, 1997; Nyhus & Curran, 2010) and the decision making literature (Frank et al., 2015; Pinner & Cavanagh, 2017; Senftleben & Scherbaum, 2021) suggest that theta (4-8 Hz activity in EEG) supports both memory and decision making processes. However, the extent to which theta dissociations in a memory task reflect decision making processes is unclear, likely due to limited investigation of EEG signals *following* a memory response. The aim of this study is to investigate the neural correlates leading up to and following a memory response.

It is well established that successful remembering is characterized by electrophyisological changes around 300 to 800 ms following stimulus onset during a memory test (Friedman & Johnson Jr., 2000; Rugg & Wilding, 2000; Voss & Paller, 2008; Addante, Ranganath, & Yonelinas, 2012). Specifically, two event-related potentials (ERPs) distinguish successful remembering of a target or studied item (hit) from successful rejection of a lure or non-studied item (correct rejection, CR). The FN400, a negative going frontal ERP component thought to reflect familiarity (Rugg et al., 1998; Mecklinger, 2000; Curran & Hancock, 2007) is more negative for CRs than hits around 300 to 500 ms after stimulus onset (Curran, 2000; Curran & Cleary, 2003; Curran, 2004) and the LPC, a late positive going parietal ERP component thought to reflect recollection (Friedman & Johnson Jr., 2000; Mecklinger, 2000) is more positive for hits than CRs around 400 to 800 ms after stimulus onset (Curran, 2000; Friedman & Johnson Jr., 2000; Curran & Cleary, 2003; Curran, 2004). Similarly, theta power is greater for hits compared to CRs, most often around 500 to 1000 ms after stimulus onset (Burgess & Gruzelier, 1997; Klimesch, Doppelmayr, Schwaiger, Winkler, & Gruber, 2000; Düzel et al., 2003). According to the drift diffusion model, recognition memory is supported by a mechanism whereby evidence accumulates over time until a threshold is reached and a decision is made (Ratcliff & McKoon, 2008). Given evidence that theta power is positively correlated with reaction times, such that theta power increases until a response is made (Jacobs, Hwang, Curran, & Kahana, 2006), theta power prior to a response may reflect evidence accumulation and/or reinstatement (Nyhus & Curran, 2010; Herweg, Solomon, & Kahana, 2020; Kota, Rugg, & Lega, 2020; Guan, Ma, Chen, Luo, & He, 2023). However, these effects are related to representation access prior to a response and as such, do not elucidate the processes that may unfold after a memory response is made.

Parallel findings from the decision making and cognitive control literature have revealed that both ERPs and theta power track errors and negative feedback signals prior to and following a response (Luu, Tucker, & Makeig, 2004; Trujillo & Allen, 2007; Cavanagh, Frank, Klein, & Allen, 2010; Cavanagh & Frank, 2014; Luft, 2014). Specifically, the error-related negativity (ERN), a negative going fronto-central ERP component that reflects decision conflict or error monitoring (Frank, Woroch, & Curran, 2005; L. Wang, Gu, Zhao, & Chen, 2020), is more negative following incorrect compared to correct responses (Gehring, Goss, Coles, Meyer, & Donchin, 1993; Cavanagh, Zambrano-Vazquez, & Allen, 2012). Similarly, across cognitive control tasks such as the Stroop task, the flanker task, and go/no-go tasks, theta power is greater following incorrect compared to correct responses and following negative relative to positive outcomes (Mazaheri, Nieuwenhuis, Dijk, & Jensen, 2009; Cohen, 2014; Cavanagh & Frank, 2014). Together, these findings suggest that post-response theta power may reflect a feedback signal or monitoring process.

Although there is evidence for post-retrieval monitoring during recognition memory (Rugg, Henson, & Robb, 2003; Hill, Horne, Koen, & Rugg, 2021), these signals are often measured following access of the representation, but prior to the memory response itself. Further complicating interpretation is that reaction times are generally faster for hits than CRs (Merkow, Burke, & Kahana, 2015; Weidemann & Kahana, 2016), meaning that stimulus-locked hit vs. CR dissociations may include both pre- and post-response related processes. There is ERP evidence that post-retrieval monitoring processes are supported by a late old/new effect over right frontal cortex (Wilding & Rugg, 1996; Johansson & Mecklinger, 2003), characterized by a positive voltage deflection that is greater for hits than CRs (Hayama, Johnson, & Rugg, 2008). However, the majority of such post-retrieval monitoring signals occur after the putative representation has been accessed, but before a behavioral response is made (Woodruff, Uncapher, & Rugg, 2006; Cruse & Wilding, 2009, 2011), leaving open the question of whether decision making mechanisms are engaged following a memory response. Greater conflict between memory decisions in a recognition task – created via differential payoff rates for correct old vs. new responses – leads to a greater post-response ERN (Curran, DeBuse, & Leynes, 2007), suggesting that control or monitoring processes may be engaged after a memory response is made.

Our hypothesis is that distinct processes occur prior to and following a memory response. To test our hypothesis, we conducted a human scalp EEG recognition memory study in which we specifically assessed response-locked theta power. By investigating response-locked signals, we can separately assess pre- and post-response related processing during both hits and CRs. We expected to replicate prior work and find greater theta power for hits compared to CRs preceding the response (equivalent to the established stimulus-locked effects, e.g. Burgess & Gruzelier, 1997; Düzel et al., 2003; Nyhus & Curran, 2010). To the extent that distinct, and potentially decision making related, processes are engaged following a memory response, we expected to find a post-response theta pattern that differed from the pre-response hit vs. CR effect. First, if there are neither memory nor decision making related signals following a memory response, theta power following both hits and CRs should return to baseline. Alternatively, because both hits and CRs are correct responses, theta power may be similarly decreased for both response types following a memory response. Finally, to the extent that successful retrieval is intrinsically rewarding (Satterthwaite et al., 2012; Speer, Bhanji, & Delgado, 2014), theta power may dissociate post-response hits from CRs such that theta power would be greater for CRs compared to hits following the memory response, reflecting a positive feedback signal for hits.

## 2 Materials and Methods

### 2.1 Participants

Forty (30 female; age range = 18-42, mean age = 21.9 years) native English speakers from the University of Virginia community participated. Our sample size of N = 40 was selected based on prior scalp EEG studies conducted in our lab (Smith, Moore, & Long, 2022; Moore & Long, 2022). All participants had normal or corrected-to-normal vision. Informed consent was obtained in accordance with University of Virginia Institutional Review Board for Social and Behavioral Research and participants were compensated for their participation. Two participants were excluded from the final dataset: one for technical difficulties that resulted in a subset of test items being presented twice during the test phase and one who failed to comply with task instructions. Thus data are reported for the remaining 38 participants. All raw, de-identified data and the associated experimental and analysis codes used in this study will be made available via the Long Term Memory Lab Website upon publication.

### 2.2 Recognition Task Experimental Design

Stimuli consisted of 1602 words, drawn from the Toronto Noun Pool (Friendly, Franklin, Hoffman, & Rubin, 1982). From this set, 288 words were randomly selected for each participant. Of these words, 192 were presented in both the study and test phase while the remaining 96 served as lures in the test phase.

#### Study Phase

In each of 12 runs, participants viewed a list containing 16 words, yielding a total of 192 trials. During each trial, participants saw a single word presented for 2000 ms followed by a 1000 ms inter-stimulus interval (ISI; Figure 1A). Participants were instructed to study the presented word in anticipation for a later memory test and did not make any behavioral responses. Each list was split evenly into two parts containing 8 words (“first associates” and “second associates,” respectively) separated by a brief 2000 ms delay. Half of the first and second associates were strongly semantically associated, as determined based on Word Association Space values (Nelson, Zhang, & McKinney, 2001) and half of the first and second associates were weakly semantically associated (Long & Kahana, 2017). Both strong and weak semantic associates were weakly semantically associated to all other study words.

**Figure 1.**
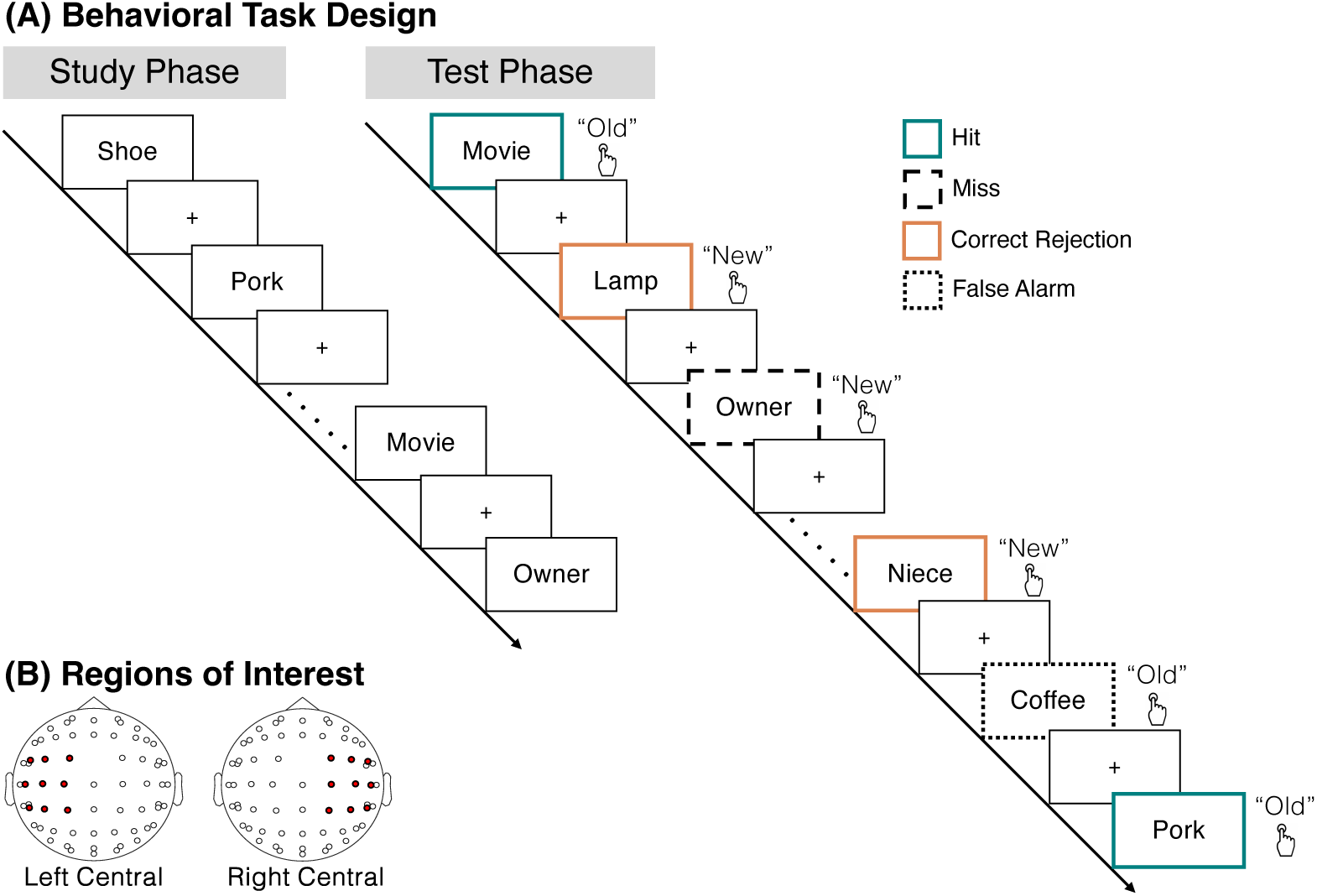
Task design. **(A)** During the study phase, participants studied individual words in anticipation of a later memory test and made no behavioral responses. After 12 runs of 16 item word lists, participants completed a recognition test phase. On each trial, participants saw either a target, a word that was presented during the study phase, or a lure, a word that was not presented during the study phase. Participants’ task was to make an old or new judgement for each word. There were one of four possible response types, hits (teal; an ‘old’ response to a target), correct rejections (orange; a ‘new’ response to a lure), misses (dashed black lines; a ‘new’ response to a target), and false alarms (dotted black lines; an ‘old’ response to a lure). Lines and colors around the boxes are shown for illustrative purposes and were not present during the actual experiment. **(B)** We analyzed two regions of interest (ROIs), a left central ROI (FC5, FC1, C3, CP5, CP1, FC3, C1, C5, CP3) and a right central ROI (CP6, CP2, C4, FC6, FC2, CP4, C6, C2, FC4).

#### Test Phase

Following the 12 study runs, participants completed the recognition test phase. On each trial, participants viewed a word which had either been presented during the study phase (target) or had not been presented (lure; Figure 1A). Participants’ task was to make an old or new judgment for each word by pressing one of two buttons (“d” or “k”). Response mappings were counterbalanced across participants. Test trials were self-paced and separated by a 1000 ms ISI. There were a total of 288 test trials with all 192 study words presented along with 96 novel lures, half of which were semantically associated to a study word. As key findings are unchanged when accounting for semantic associations, we do not consider them further.

### 2.3 EEG Data Acquisition and Preprocessing

All acquisition and preprocessing methods are based on our previous work (Smith et al., 2022); for clarity we use the same text as previously reported. EEG recordings were collected using a BrainVision system and an ActiCap equipped with 64 Ag/AgCl active electrodes positioned according to the extended 10-20 system. All electrodes were digitized at a sampling rate of 1000 Hz and were referenced to electrode FCz. Offline, electrodes were later converted to an average reference. Impedances of all electrodes were kept below 50kΩ. Electrodes that demonstrated high impedance or poor contact with the scalp were excluded from the average reference. Bad electrodes were determined by voltage thresholding (see below).

Custom python codes were used to process the EEG data. We applied a high pass filter at 0.1 Hz, followed by a notch filter at 60 Hz and harmonics of 60 Hz to each participant’s raw EEG data. We then performed three preprocessing steps (Nolan, Whelan, & Reilly, 2010) to identify electrodes with severe artifacts. First, we calculated the mean correlation between each electrode and all other electrodes as electrodes should be moderately correlated with other electrodes due to volume conduction. We z-scored these means across electrodes and rejected electrodes with z-scores less than -3. Second, we calculated the variance for each electrode, as electrodes with very high or low variance across a session are likely dominated by noise or have poor contact with the scalp. We then z-scored variance across electrodes and rejected electrodes with a *|*z*| >* = 3. Finally, we expect many electrical signals to be autocorrelated, but signals generated by the brain versus noise are likely to have different forms of autocorrelation. Therefore, we calculated the Hurst exponent, a measure of long-range autocorrelation, for each electrode and rejected electrodes with a *|*z*| >* = 3. Electrodes marked as bad by this procedure were excluded from the average re-reference. We then calculated the average voltage across all remaining electrodes at each time sample and re-referenced the data by subtracting the average voltage from the filtered EEG data. We used wavelet-enhanced independent component analysis (Castellanos & Makarov, 2006) to remove artifacts from eyeblinks and saccades.

### 2.4 EEG Data Analysis

We applied the Morlet wavelet transform (wave number 6) to the entire EEG time series across electrodes, for each of 46 logarithmically spaced frequencies (2-100 Hz; Long & Kahana, 2015). Because we hypothesized distinct processes occur prior to and following a memory response, after log-transforming the power we focused exclusively on test-phase data. We then downsampled the test-phase data by taking a moving average across 100 ms time intervals from -1000 to 3000 ms relative to the response and sliding the window every 25 ms, resulting in 157 time intervals (40 non-overlapping). Mean and standard deviation power were calculated across all trials and across time points for each frequency. Power values were then z-transformed by subtracting the mean and dividing by the standard deviation power. We focus exclusively on the theta band (4-8 Hz) for all analyses.

### 2.5 Regions of Interest

We examined theta power across two regions of interest (ROIs; Figure 1B), left central (FC5, FC1, C3, CP5, CP1, FC3, C1, C5, CP3) and right central (CP6, CP2, C4, FC6, FC2, CP4, C6, C2, FC4) based on prior work in which theta power in these regions dissociates hits and correct rejections (Nyhus & Curran, 2010).

### 2.6 Univariate Analyses

To test the effect of response type on theta power leading up to and following memory responses, our two conditions of interest were hits (correctly recognized targets) and correct rejections (CRs, correctly rejected lures). We compared theta power across hits and CRs separately for each ROI. For each participant, we calculated z-transformed theta power across both ROIs in each of the two conditions, across 100 ms time intervals from 500 ms pre-response to 1000 ms post-response. For a direct comparison of pre- response and post-response theta power, we further averaged z-transformed theta power over the 500 ms pre-response and 500 ms post-response interval separately for hits and CRs. We selected 500 ms as our pre-response interval based on our prior work investigating contextually mediated retrieval processes in the hippocampus (Long et al., 2017).

### 2.7 Statistical Analyses

We used a repeated measures ANOVA (rmANOVA) to assess the distribution of reaction times (RTs) for hits and CRs. For post hoc comparisons across RTs, we used false discovery rate (FDR; *p* = .05) correction (Benjamini & Hochberg, 1995) to correct for multiple comparisons. We used rmANOVAs and paired-sample *t* -tests to assess the effect of response type (hits, CRs) and time interval (pre-response, post-response) on theta power.

## 3 Results

Our first goal was to measure memory discrimination (d’) to ensure that participants were following directions and able to discriminate between targets and lures. For each participant, we calculated d’ by subtracting the normalized false alarm rate (the percentage of lures that were incorrectly identified as ‘old’) from the normalized hit rate (the percentage of targets that were correctly identified as ‘old’). The average d’ was 1.75 (SD = 0.58; Figure 2A), indicating that participants were able to successfully distinguish targets from lures.

**Figure 2.**
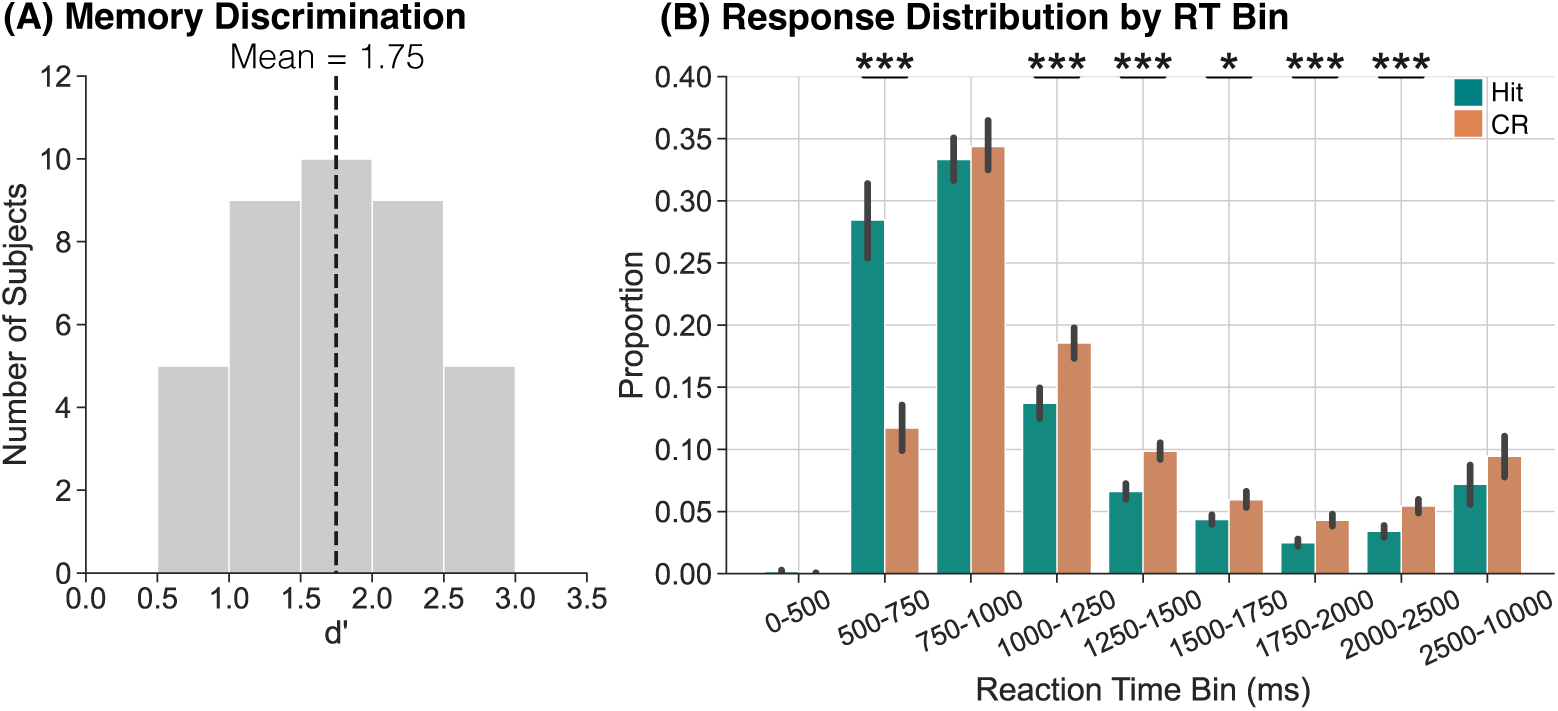
Memory discrimination and responses as a function of reaction time bin. **(A)** We used d’ to assess memory discrimination. Participants were able to correctly discriminate between targets and lures. **(B)** We assessed the proportion of hits (teal) and CRs (orange) as a function of RT bin. The highest proportion of hits and CRs occurs in the 750-1000 ms bin and a significantly greater proportion of hits compared to CRs occur within the 500-750 ms bin. Error bars reflect standard error of the mean. *p *<* 0.05; ***p *<*.001 (FDR-corrected).

Having found that participants are able to discriminate targets and lures, we next assessed median reaction times (RTs) for hits and CRs. To the extent that RTs reliably differ between hits and CRs, stimulus- locked neural dissociations between these conditions may be driven by engagement of different processes, e.g. memory vs. decision making, rather than differential engagement of the same process. That is, if hits occur more quickly than CRs, a stimulus-locked comparison between the two conditions could reflect a comparison between post-response processes for hits vs. pre-response processes for CRs. The average median RT for hits (M = 954.9, SD = 419.6) was significantly faster than for CRs (M = 1122.8, SD = 317.3, *t* _37_ = -4.427, *p* = 0.0001, *d* = 0.4514). This RT difference suggests that stimulus-locked theta dissociations could be driven by dissociations in pre- vs. post-response processes across hits and CRs.

Given the dissociation in median RT between hits and CRs, we next sought to compare the distribution of RTs across conditions to determine when relative to stimulus onset the majority of hits and CRs occur. We grouped RTs into nine bins selected to cover the full range of RTs with higher resolution in the faster (*<* 2500 ms) bins: 0-500 ms, 500-750 ms, 750-1000 ms, 1000-1250 ms, 1250-1500 ms, 1500-1750 ms, 1750-2000 ms, 2000-2500 ms, and 2500-10000 ms (Figure 2B). We calculated the proportion of hits and CRs that occurred within each RT bin for each participant and then averaged those proportions across all participants. We conducted a 2 *×* 9 rmANOVA with factors of response type (hit, CR) and RT bin. We do not find a significant main effect of response type (*F* _1,37_ = 0.51, *p* = 0.479, *η_p_*^2^ = 0.01). We find a significant main effect of RT bin (*F* _8,296_ = 74.48, *p <* 0.0001, *η_p_*^2^ = 0.67) and a significant interaction between response type and RT bin (*F* _8,296_ = 22.36, *p <* 0.0001, *η_p_*^2^ = 0.38). We report the results of post hoc *t* -tests comparing proportions of hits and CRs within each RT bin in Table 1 and highlight the key findings below. We find that the highest proportion of responses occurs in the 750-1000 ms RT bin for both hits and CRs. However, a significantly greater proportion of hits (M = 0.28, SD = 0.19) occur within the 500-750 ms RT bin compared to CRs (M = 0.12, SD = 0.11). Given that a larger proportion of hits occur more quickly compared to CRs, neural dissociations observed within the 500-750 ms interval may reflect a difference in post-response hit-related processing and pre-response CR-related processing. To the extent that decision making mechanisms are engaged following a memory response, the stimulus-locked contrast between hits and CRs may reflect a comparison between memory and decision making processes.

**Table 1.** Post hoc *t* -tests comparing the proportion of hits and CRs in each RT bin.

| RT Bin (ms) | Hits |  | CRs |  | Hits vs CRs |  |  |
| --- | --- | --- | --- | --- | --- | --- | --- |
| | Mean | SD | Mean | SD | $t_{37}$ | $p$ | $d$ |
| 0-500 | 0.002 | 0.006 | 0.0004 | 0.002 | 1.579 | 0.1229 | 0.3252 |
| 500-750 | 0.28 | 0.19 | 0.12 | 0.11 | 7.058 | **< 0.0001** | 1.081 |
| 750-1000 | 0.33 | 0.11 | 0.34 | 0.13 | -0.466 | 0.6438 | 0.0898 |
| 1000-1250 | 0.14 | 0.08 | 0.19 | 0.07 | -4.198 | **0.0002** | 0.6536 |
| 1250-1500 | 0.07 | 0.04 | 0.10 | 0.04 | -4.425 | **0.0001** | 0.7708 |
| 1500-1750 | 0.04 | 0.02 | 0.06 | 0.04 | -2.549 | **0.02** | 0.4681 |
| 1750-2000 | 0.02 | 0.02 | 0.04 | 0.03 | -4.012 | **0.0003** | 0.7581 |
| 2000-2500 | 0.03 | 0.03 | 0.05 | 0.03 | -3.973 | **0.0003** | 0.6373 |
| 2500-10000 | 0.07 | 0.10 | 0.09 | 0.10 | -2.009 | 0.052 | 0.2192 |

*Note:* bold values indicate tests that survive FDR correction.

Our hypothesis is that distinct memory and decision making processes occur preceding and following a memory response, meaning that we should find differential theta power engagement pre- and post-response. Specifically, we should replicate past findings of greater theta power for hits compared to CRs pre-response, reflecting memory related processing. To the extent that decision making processes are engaged following a response, we should either find decreased theta power for both hits and CRs – as both conditions are correct responses – or we may find a theta dissociation between hits and CRs. That is, to the extent that successful retrieval is intrinsically rewarding, post-response theta power should be decreased for hits compared to CRs. We conducted a 2 *×* 2 *×* 15 rmANOVA with factors of response type (hit, CR), ROI (left central; LC, right central; RC) and time interval (-500 to 1000 ms in fifteen 100 ms intervals). We find a significant main effect of ROI (*F* _1,37_ = 11.36, *p* = 0.002, *η_p_*^2^ = 0.23) driven by greater theta power for the RC compared to the LC ROI. We find a significant main effect of time interval (*F* _14,518_ = 21.59, *p <* 0.0001, *η_p_*^2^ = 0.37). We do not find a significant main effect of response type (*F* _1,37_ = 0.172, *p* = 0.681, *η_p_*^2^ = 0.005) or interaction between response type and ROI (*F* _1,37_ = 0.218, *p* = 0.643, *η_p_*^2^ = 0.006). We find a significant interaction between ROI and time interval (*F* _14,518_ = 3.408, *p <* 0.0001, *η_p_*^2^ = 0.08) and between response type and time interval (*F* _14,518_ = 6.127, *p <* 0.0001, *η_p_*^2^ = 0.14). We find a significant three-way interaction between response type, ROI and time interval (*F* _14,518_ = 2.23, *p* = 0.006, *η_p_*^2^ = 0.06).

Given the significant three-way interaction between response type, ROI, and time interval, we next performed follow-up post-hoc ANOVAs over time separately for each ROI. We conducted two 2 *×* 15 rmANOVAs with factors of response type (hit, CR) and time interval (-500 to 1000 ms in fifteen 100 ms intervals). For both ROIs (Figure 3), we find a significant main effect of time interval (LC: *F* _14,518_ = 19.76, *p <* 0.0001, *η_p_*^2^ = 0.35; RC: *F* _14,518_ = 20.91, *p <* 0.0001, *η_p_*^2^ = 0.36), no main effect of response type (LC: *F* _1,37_ = 0.382, *p* = 0.54, *η_p_*^2^ = 0.01; RC: *F* _1,37_ = 0.007, *p* = 0.934, *η_p_*^2^ = 0.0002) and a significant interaction between response type and time interval (LC: *F* _14,518_ = 5.373, *p <* 0.0001, *η_p_*^2^ = 0.13; RC: *F* _14,518_ = 3.791, *p <* 0.0001, *η_p_*^2^ = 0.09). These results demonstrate that theta power dissociations between hits and CRs vary across time interval, suggesting that differential processes may be engaged pre- vs. post-response that distinguish these response types.

**Figure 3.**
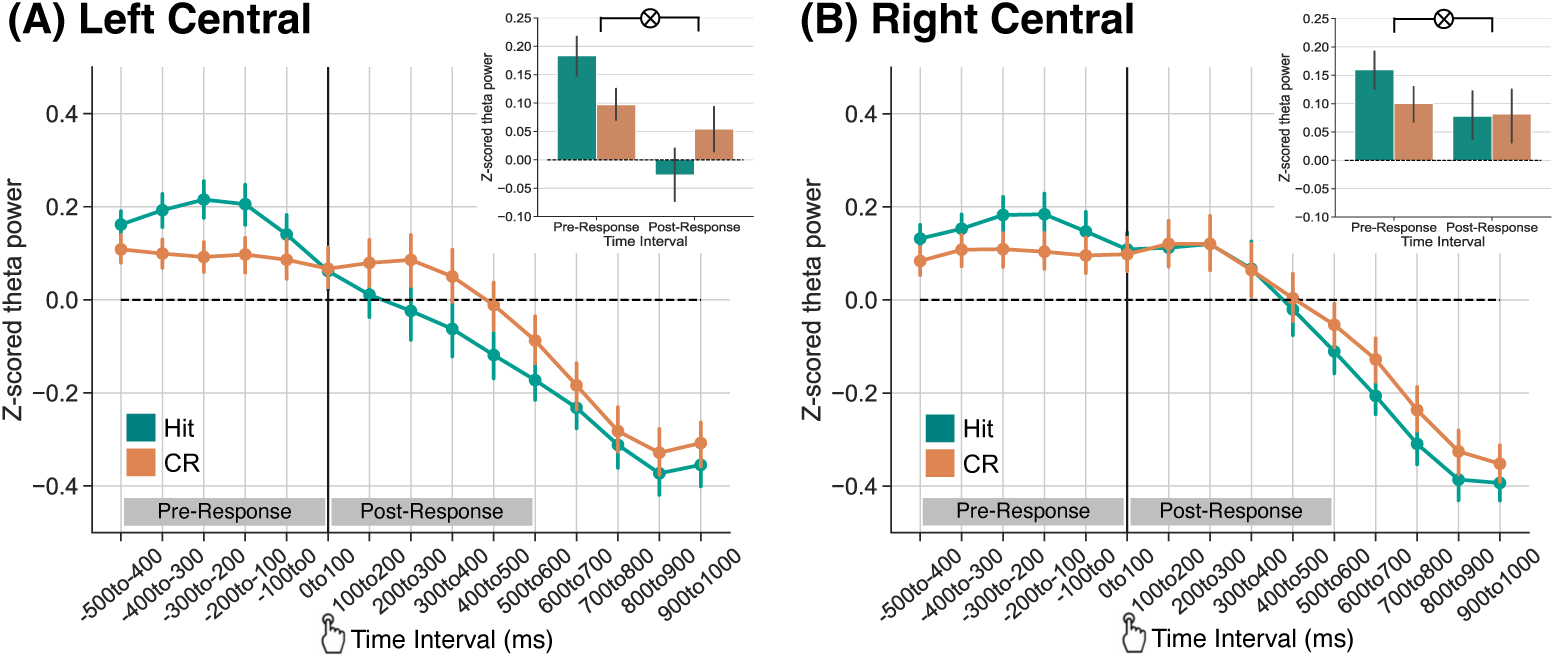
Theta power dissociations across hits and correct rejections preceding and following memory responses. Response-locked z-transformed theta power (4-8 Hz) for the left and right central ROIs. The solid vertical black line indicates when the response was made. Hits are shown in teal, correct rejections (CRs) are shown in orange. Error bars reflect standard error of the mean. **(A)** Over the left central ROI, we find a significant interaction between response type and time interval (p *<* 0.001) driven by greater pre-response theta power for hits than CRs and numerically greater post-response theta power for CRs than hits. **(B)** Over the right central ROI, we find a significant interaction between response type and time interval (p = 0.027) driven by greater theta power for hits than CRs during the pre-response time interval.

To specifically test for a pre- vs. post-response dissociation in theta power for hits and CRs, we averaged signals within the 500 ms pre- and post-response time intervals (Figure 3, insets). We conducted two 2 *×* 2 rmANOVAs, one for each ROI, with factors of response type (hit, CR) and time interval (pre-response, post-response). In LC, we find a significant main effect of time interval (*F* _1,37_ = 5.832, *p* = 0.0208, *η_p_*^2^ = 0.14) driven by greater pre-response theta power compared to post-response theta power. We do not find a significant main effect of response type (*F* _1,37_ = 0.009, *p* = 0.925, *η_p_*^2^ = 0.0002). We find a significant interaction between response type and time interval (*F* _1,37_ = 16.18, *p* = 0.0003, *η_p_*^2^ = 0.30). This interaction was driven by greater theta power for hits (M = 0.18, SD = 0.21) relative to CRs (M = 0.10, SD = 0.18) in the pre-response time interval (*t* _37_ = 2.559, *p* = 0.0147, *d* = 0.4459) and numerically greater theta power for CRs (M = 0.05, SD = 0.25) relative to hits (M = -0.03, SD = 0.29) in the post-response time interval (*t* _37_ = 2.021, *p* = 0.0506, *d* = 0.3019). In RC, we do not find a significant main effect of time interval (*F* _1,37_ = 0.708, *p* = 0.406, *η_p_*^2^ = 0.02) or response type (*F* _1,37_ = 0.871, *p* = 0.357, *η_p_*^2^ = 0.02). We find a significant interaction between response type and time interval (*F* _1,37_ = 5.306, *p* = 0.027, *η_p_*^2^ = 0.13). This interaction was driven by numerically greater theta power for hits (M = 0.16, SD = 0.21) relative to CRs (M = 0.10, SD = 0.19) in the pre-response time interval (*t* _37_ = 1.911, *p* = 0.0637, *d* = 0.295) and no difference in theta power for hits (M = 0.08, SD = 0.24) relative to CRs (M = 0.08, SD = 0.29) in the post-response time interval (*t* _37_ = -0.1160, *p* = 0.9083, *d* = 0.015). Together, these results indicate that theta power is modulated by successful retrieval leading up to and following memory responses.

Prior work (Herweg et al., 2020; Guan et al., 2023) suggests that the pre-response theta power dissociation that we observe specifically reflects evidence accumulation or reinstatement that ultimately supports recollection. The post-response theta power effects may likewise reflect a feedback signal in response to recollected content. Due to the current task design we cannot directly measure recollection and familiarity; however, we can leverage reaction time (RT) as a coarse assay of confidence. The general assumption is that compared to trials with slow RTs, trials with fast RTs reflect faster evidence accumulation (Mulder & van Maanen, 2013; Shenhav, Straccia, Musslick, Cohen, & Botvinick, 2018; Rollwage et al., 2020) and greater confidence (Ratcliff, 1978; Ratcliff & Starns, 2009; Weidemann & Kahana, 2016). To divide the trials based on RT, we calculated the median RT across all correct responses (hits and CRs) for each participant, and labeled hits as either ‘fast’ (those below the median RT) or ‘slow’ (those above the median RT). To the extent that fast hits reflect strongly remembered experiences or the degree of evidence accumulation, we should find differential theta power engagement pre- and post-response. Specifically, we should find greater pre-response theta power for fast hits compared to CRs, as there is no experience to remember or reinstate during a CR. To the extent that post-response theta power reflects a feedback signal based on reinstated content, we should find decreased theta power for fast hits compared to CRs.

To specifically test for a pre- vs. post-response dissociation in theta power for fast hits, slow hits, and CRs, we averaged signals within the 500 ms pre- and post-response time intervals (Figure 4). We conducted two 2 *×* 3 rmANOVAs, one for each ROI, with factors of time interval (pre-response, post-response) and response type (fast hit, slow hit, CR). In LC, we find a significant main effect of time interval (*F* _1,37_ = 7.459, *p* = 0.0096, *η_p_*^2^ = 0.17) driven by greater pre-response theta power compared to post-response theta power. We do not find a significant main effect of response type (*F* _2,74_ = 0.019, *p* = 0.981, *η_p_*^2^ = 0.0005). We find a significant interaction between response type and time interval (*F* _2,74_ = 14.93, *p <* 0.0001, *η_p_*^2^ = 0.29). This interaction was driven by a significant interaction between fast hits and CRs (*F* _1,37_ = 24.56, *p <* 0.0001, *η_p_*^2^ = 0.40), whereby theta power was significantly greater for fast hits (M = 0.22, SD = 0.23) relative to CRs (M = 0.10, SD = 0.18) in the pre-response time interval (*t* _37_ = 3.080, *p* = 0.0039, *d* = 0.5789) and for CRs (M = 0.05, SD = 0.25) relative to fast hits (M = -0.07, SD = 0.33) in the post-response time interval (*t* _37_ = 2.554, *p* = 0.0149, *d* = 0.4161). The interaction between slow hits and CRs was not significant (*F* _1,37_ = 2.892, *p* = 0.0974, *η_p_*^2^ = 0.07).

**Figure 4.**
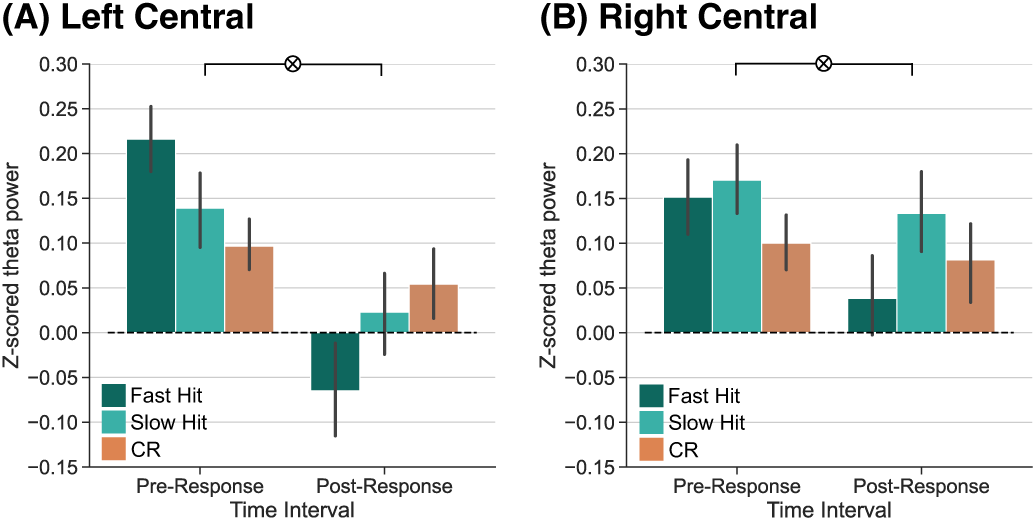
Theta power dissociations across fast hits, slow hits, and correct rejections preceding and following memory responses. Response-locked z-transformed theta power (4-8 Hz) for the left and right central ROIs. Fast hits are shown in dark teal, slow hits are shown in light teal and correct rejections (CRs) are shown in orange. Error bars reflect standard error of the mean. **(A)** Over the left central ROI, we find a significant interaction between response type and time interval (*p <* 0.0001) driven by a significant pre-post interaction between fast hits and CRs (*p <* 0.0001). **(B)** Over the right central ROI, we find a significant interaction between response type and time interval (*p* = 0.0206) driven by a significant pre-post interaction between fast hits and CRs (*p* = 0.008).

In RC, we do not find a significant main effect of time interval (*F*_1,37_ = 0.87, *p* = 0.357, *η_p_*^2^ = 0.02) or response type (*F* _2,74_ = 1.938, *p* = 0.151, *η_p_*^2^ = 0.05). We find a significant interaction between response type and time interval (*F* _2,74_ = 4.095, *p* = 0.0206, *η_p_*^2^ = 0.10). This interaction was driven by a significant interaction between fast hits and CRs (*F* _1,37_ = 7.885, *p* = 0.008, *η_p_*^2^ = 0.18), whereby theta power was numerically greater for fast hits (M = 0.15, SD = 0.26) relative to CRs (M = 0.10, SD = 0.19) in the pre-response time interval (*t* _37_ = 1.274, *p* = 0.2107, *d* = 0.2278) and numerically greater for CRs (M = 0.08, SD = 0.29) relative to fast hits (M = 0.04, SD = 0.27) in the post-response time interval (*t* _37_ = 0.9688, *p* = 0.3389, *d* = 0.1549). The interaction between slow hits and CRs was not significant (*F* _1,37_ = 0.306, *p* = 0.583, *η_p_*^2^ = 0.008). Together, these results suggest that post-response theta power dissociations may reflect a positive feedback signal in response to strongly remembered experiences.

## 4 Discussion

The goal of the current study was to test the hypothesis that distinct processes occur prior to and following a memory response. We recorded scalp EEG while participants performed a recognition memory task. Crucially, we focused our analyses on response-locked test-phase data to separate the processes occurring before and after a memory response. We show that reaction times (RTs) are faster for hits compared to correct rejections (CRs) and that a greater proportion of hits occur within 500-750 ms of stimulus onset compared to CRs. We replicate established findings (Nyhus & Curran, 2010) that preceding a memory response, theta power (4-8Hz) in central electrodes is greater for hits compared to CRs. We show that these pre-response theta power dissociations ‘flip’ in left central electrodes following the memory response. We find that this post-response ‘flip’ is specific to hits committed quickly, potentially reflecting a positive feedback signal for strongly remembered experiences. Together, these findings suggest that there are potentially distinct memory and decision making processes engaged preceding and following a memory response that are modulated by successful retrieval.

We find faster RTs for hits compared to CRs, replicating past findings (Merkow et al., 2015; Weidemann & Kahana, 2016). Faster RTs for hits may be driven by greater memory strength (Verde & Rotello, 2007; Wixted, 2007) and/or greater contextual reinstatement (Gordon, Rissman, Kiani, & Wagner, 2014; Hanczakowski, Zawadzka, & Macken, 2015). However, these RT differences indicate that traditional hit vs. CR comparisons of stimulus-locked data may capture both pre- and post-response related processes. We specifically find that a larger proportion of hits occur within 500-750 ms of stimulus onset compared to CRs meaning that estimates of neural signals within this time window may reveal differences in post-response processes related to successful retrieval vs. pre-response processes related to correctly rejecting lures. Thus, our results highlight the importance of utilizing response-locked data to investigate the distinct processes leading up to and following a memory response.

Consistent with prior EEG work (Burgess & Gruzelier, 1997; Klimesch et al., 1997, 2000; Düzel et al., 2003; Nyhus & Curran, 2010), we find greater pre-response theta power for hits than CRs. EEG power in the theta frequency band has been proposed to coordinate cortical areas, supporting the ability to encode and retrieve contextual information about time and space that is central to episodic memory (Hasselmo & Stern, 2014). Successful memory retrieval depends on the access of an internal representation of a past experience. This access may take the form of reinstatement, wherein encoded content is reconstructed during retrieval (Danker & Anderson, 2010), which may specifically be supported by coordination between cortical areas (Herweg et al., 2020). The pre-response theta power dissociations that we observe in the current study may reflect evidence accumulation, as in the drift diffusion model (Ratcliff & McKoon, 2008; Ratcliff, Smith, Brown, & McKoon, 2016). This interpretation is consistent with our finding of robust pre-response theta power dissociations specifically between CRs and fast hits. Fast RTs are thought to reflect rapid evidence accumulation (Mulder & van Maanen, 2013; Shenhav et al., 2018; Rollwage et al., 2020), thus greater pre-response theta power for fast hits compared to CRs may reflect faster evidence accumulation for strongly remembered experiences.

Following a memory response, we find a ‘flip’ in theta power over the left central ROI, such that theta power is greater for CRs than hits. Although prior work has demonstrated that post-retrieval monitoring processes are engaged following memory retrieval (Rugg et al., 2003; Hill et al., 2021), the majority of these findings reflect signals that precede a behavioral response. Our interpretation is that the post- response theta power dissociation between hits and CRs may reflect a feedback signal. This interpretation is consistent with work from the cognitive control literature showing that theta power increases for incorrect compared to correct responses and following negative relative to positive outcomes (Cavanagh et al., 2010; Cavanagh & Frank, 2014). Prior work has proposed that frontal midline theta (FMT) is associated with reward processing – specifically that the FMT is larger following negative feedback or monetary loss (Cohen, Elger, & Ranganath, 2007; Marco-Pallarés et al., 2008) – and is a mechanism for communication between brain regions (Glazer, Kelley, Pornpattananangkul, Mittal, & Nusslock, 2018). As both hits and CRs constitute correct responses, we may have anticipated decreased post-response theta power for both response types. However, only hits reflect successful retrieval. Thus, the post-response decrease in theta power for hits may reflect positive feedback specifically in response to successful retrieval. We specifically find that the pre- vs. post-response dissociation is specific to fast hits. To the extent that fast hits reflect highly confident responses (Ratcliff, 1978; Ratcliff & Starns, 2009; Weidemann & Kahana, 2016), the post-response theta power decrease for fast hits may reflect a positive feedback signal. Successful retrieval may be intrinsically rewarding (Satterthwaite et al., 2012; Speer et al., 2014) and neuro-imaging work has repeatedly shown that reward-related regions (e.g. striatum) are more active during hits compared to CRs (Achim & Lepage, 2005; de Zubicaray, McMahon, Eastburn, Finnigan, & Humphreys, 2005; Henson, Hornberger, & Rugg, 2005; Fliessbach, Weis, Klaver, Elger, & Weber, 2006; Spaniol et al., 2009; Schwarze, Bingel, Badre, & Sommer, 2013; Clos, Schwarze, Gluth, Bunzeck, & Sommer, 2015). Taken together, the post-response theta dissociation may represent a positive feedback signal in response to successful retrieval. The direct investigation of post-response feedback signals during memory tests presents an exciting avenue for future work.

A critical open question is how the observed RT distributions and pre- vs. post-response theta dissociations relate to the established processes of recollection and familiarity. Recollection is the retrieval of contextual details (Yonelinas, 2001a; Diana, Vilberg, & Reder, 2005; Gimbel & Brewer, 2011; Addante et al., 2012) and familiarity is memory strength without detailed retrieval (Yonelinas, 2001a; Diana et al., 2005; Gimbel & Brewer, 2011; Addante et al., 2012). There is mixed evidence as regards RTs for recollection and familiarity, with some evidence that recollection responses are faster than familiarity responses (Diana et al., 2005; Gimbel & Brewer, 2011; Herweg et al., 2016) and some evidence that recollection responses are slower than familiarity responses (Atkinson & Juola, 1974; Jacoby, 1991; Besson, Ceccaldi, Didic, & Barbeau, 2012). Due to our task design, we cannot disambiguate recollection from familiarity based responses, but we can leverage RT as a coarse assay of confidence. Our assumption is that compared to slow hits, fast hits reflect higher confident responses (Ratcliff, 1978; Ratcliff & Starns, 2009; Weidemann & Kahana, 2016). High confident responses may be supported by the recollection of contextual details (Yonelinas, 2001a, 2001b), in which case elevated pre-response theta power may reflect the reinstatement of contextual details during fast hits. Likewise, the decreased post-response theta power following fast hits may reflect a positive feedback signal in response specifically to recollected experiences. Future work will be needed to directly test this possibility.

Although we did not anticipate hemispheric differences, we consistently found influences of ROI on theta power. Overall, the effects that we observe are numerically stronger in the left central, relative to right central, ROI. This hemispheric asymmetry may be driven by the stimuli used and/or intrinsic hemispheric connections (D. Wang, Buckner, & Liu, 2014). We used visually presented words in the current study and verbal stimuli are well known to recruit the left hemisphere (de Zubicaray, Miozzo, Johnson, Schiller, & McMahon, 2011; Vigneau et al., 2011; Price, 2012; Ries, Dronkers, & Knight, 2016), including during memory tasks (Kelley et al., 1998; Kim, 2011). Investigation of resting state data has shown that cortical networks have intrinsic within-hemisphere connections which may enable control over the specific functions or processes that are engaged (D. Wang et al., 2014).

Together, our findings suggest that distinct processes occur prior to and following a memory response and, in particular, that decision making processes may follow successful retrieval. A direction for future research will be to directly investigate the extent to which the post-response theta dissociation reflects a feedback signal. More broadly, we contribute to a growing body of literature characterizing the role of theta activity in successful memory retrieval.

## Acknowledgments

We thank Yuju Hong for assistance with data collection. This work was supported by a grant from the National Institutes of Health (NINDS R01 NS132872, PI: NML).

## Data and Code Availability

The raw, de-identified data and the associated experimental and analysis codes used in this study will be made accessible via the Long Term Memory laboratory website (https://longtermmemorylab.com) upon publication.

## Author Contributions

Devyn E. Smith: Conceptualization; Data curation; Formal analysis; Software; Supervision; Visualization; Writing—Original draft; Writing—Review & editing. Justin R. Wheelock: Formal analysis; Software; Visualization; Writing—Original draft; Writing—Review & editing. Nicole M. Long: Conceptualization; Formal analysis; Funding acquisition; Project administration; Software; Supervision; Visualization; Writing—Original draft; Writing—Review & editing.

## Declaration of Competing Interests

The authors declare no competing financial interests.

## Notes

### Competing Interest Statement

The authors have declared no competing interest.

